## Supplementary Information for "A maximum-entropy model to predict 3D structural ensembles of chromatins from pairwise distances: Applications to Interphase Chromosomes and Structural Variants"

### Supplementary Information: A method to predict 3D structural ensembles of chromatins from pair-wise distances: Applications to Interphase Chromosomes and Structural Variants

Guang Shi[1,2,*] and D. Thirumalai[1,†]

[1]*Department of Chemistry, The University of Texas at Austin, Austin, Texas 78712, USA*

[2]*Current Address: Department of Materials Science, University of Illinois, Urbana, Illinois 61801, USA*

**SUPPLEMENTARY NOTE 1**

We show that

$$P^{\text{MaxEnt}}(\{\boldsymbol{x}_i\}) = \frac{1}{Z}\text{exp}\big(-\sum_{i<j}^{N} k_{ij}||\boldsymbol{x}_i - \boldsymbol{x}_j||^2\big), \tag{1}$$

which is the the starting point in the DIMES method (see the main text), is a multivariate normal distribution. In the above equation, $\boldsymbol{x}_i = [x_{1i}, x_{2i}, x_{3i}]$ are the 3D the coordinates of the $i^{th}$ locus. We write Eq.1 in a matrix form,

$$P^{\text{MaxEnt}}(\{\boldsymbol{x}_i\}) = \frac{1}{Z}\prod_{p}\text{exp}\big(\boldsymbol{X}_p^T\boldsymbol{K}\boldsymbol{X}_p\big), \tag{2}$$

where $\boldsymbol{X}_p = [x_{p1}, x_{p2}, \cdots, x_{pN}]^T$, subscript $p \in (1,2,3)$ denote the three spatial dimensions, and $N$ is the total number of loci. The connectivity matrix, $\boldsymbol{K}$, is given by,

$$K_{ij} = k_{ij}, \text{ if } i \neq j \text{ and } K_{ii} = -\sum_{j\neq i} k_{ij}. \tag{3}$$

Eq.2 is a product of three multivariate normal distributions with covariance matrix $\boldsymbol{\Sigma} = -\boldsymbol{K}^+$ and mean 0, i.e. $\boldsymbol{X}_p \sim \mathcal{N}(\boldsymbol{0}, \boldsymbol{\Sigma})$ for $p = 1,2,3$ ($\mathcal{N}(\boldsymbol{0}, \boldsymbol{\Sigma})$ is the multivariate Gaussian distribution). $\boldsymbol{K}^+$ is the Moore–Penrose inverse (pseudoinverse) of $\boldsymbol{K}$. Note that Moore–Penrose inverse ensures that, even if $\boldsymbol{\Sigma}$ is not full rank, the distribution is properly defined. In DIMES, $\boldsymbol{\Sigma}$ has exact one zero eigenvalue, which corresponds to the zero mode (center of the system)

Note that for $P^{\text{MaxEnt}}(\{\boldsymbol{x}_i\})$ to be a normalizable probability density distribution, it is known that $\boldsymbol{\Sigma}$ has to be positive semidefinite ($\boldsymbol{\Sigma} \geq 0$). $\boldsymbol{\Sigma}$ with zero eigenvalue (not full rank) corresponds to the degenerate case, which is expected for a system that is translationally invariant. Hence, it is necessary that $\boldsymbol{K}$ be negative semidefinite. If $k_{ij} \geq 0$ for all $i, j$, as is the case for the Generalized Rouse Model [1, 2], $\boldsymbol{K}$ can be proven to be negative semidefinite. Even if $k_{ij} < 0$ for some $i, j$, $\boldsymbol{K}$ can still be negative semidefinite. This means that negative $k_{ij}$ values are allowed as long as the matrix $\boldsymbol{K}$ remains semidefinite.

*

†

#### SUPPLEMENTARY NOTE 2

The purpose of this note is to prove that there exists a unique set of $k_{ij}$s that satisfy the set of constraints, which in our problem is specified the average squared pair-wise distances, $\langle ||\boldsymbol{x}_i - \boldsymbol{x}_j||^2 \rangle = \langle r_{ij}^2 \rangle$. The constraints could be measured or calculated.

The distribution of coordinates of the loci is given by Eq.1. Following the derivation in our previous work [1, 3], we have,

$$\langle r_{ij}^2 \rangle = 3\omega_{ij}^2 \tag{4}$$

where $\omega_{ij}^2 = \Sigma_{ii} + \Sigma_{jj} - 2\Sigma_{ij}$, and $\boldsymbol{\Sigma} = -\boldsymbol{K}^+$. Note that the whole set of equations $\omega_{ij}^2 = \Sigma_{ii} + \Sigma_{jj} - 2\Sigma_{ij}$ has the same number of unknown variables ($\Sigma_{ij}$) as the number of equations, which leads to a unique solution for $\Sigma_{ij}$ given the values of $\langle r_{ij}^2 \rangle$. We may obtain $\boldsymbol{K}$ in principle using $\boldsymbol{K} = -\boldsymbol{\Sigma}^+$. Because $\boldsymbol{\Sigma}$ is unique, so is $\boldsymbol{K}$.

Note that this also shows that $\boldsymbol{K}$ can be obtained directly without any optimization procedure. However, there are three issues associated with this method. First, this involves a matrix inversion operation, which is usually numerically unstable for a large matrix. Second, if the target pair-wise distance matrix is not a proper distance matrix (for example it does not satisfy triangle inequality) then the resulting $\boldsymbol{\Sigma}$ is not positive semidefinite, which would result in an invalid distribution (Eq.2). Third, the direct calculation does not allow for regularization (see Supplementary Note 3.C), which is needed in practice.

#### SUPPLEMENTARY NOTE 3 – ALGORITHMS

##### A. Optimization algorithms

The objective in the application of the maximum entropy principle is to find a distribution that maximizes the entropy among all the distributions, which also satisfies a set of specified constraints. In a typical application, the constraints are the $m$ mean values, which are functions over certain random variables. We write these constraints as $\mathbb{E}[f_i(\boldsymbol{x})] = \hat{a}_i$ where $f_i(\boldsymbol{x})$ is the $i^{th}$ function over the random variable $\boldsymbol{x}$. The maximum entropy distribution is given by the following form,

$$P^{\text{MaxEnt}}(\boldsymbol{x}) = \frac{1}{Z}\exp\Big(\sum_i^m \theta_i f_i(\boldsymbol{x})\Big) \tag{5}$$

where $\theta_i$ are the Lagrange multipliers that have to be determined. The values of $\theta_i$ are determined so that the constraints are satisfied, that is,

$$\int \mathrm{d}\boldsymbol{x} P^{\mathrm{MaxEnt}}(\boldsymbol{x}) f_i(\boldsymbol{x}) = \hat{a}_i, \text{ for all } i \tag{6}$$

According to [4, 5], the problem of finding the values of $\theta_i$ is equivalent to minimizing the objective function,

$$L(\boldsymbol{\theta}) = \ln Z(\boldsymbol{\theta}) - \boldsymbol{\theta}^T \hat{\boldsymbol{a}} \tag{7}$$

where $\boldsymbol{\theta} = [\theta_1, \theta_2, \cdots, \theta_m]$, $\hat{\boldsymbol{a}} = [\hat{a}_1, \hat{a}_2, \cdots, \hat{a}_m]^T$, and $Z(\boldsymbol{\theta}) = \int \mathrm{d}\boldsymbol{x} \, \exp\big(\sum_i^m \theta_i f_i(\boldsymbol{x})\big)$ is the normalization factor (partition function). It can be shown that $L(\boldsymbol{\theta})$ is a global minimum when all the constraints $\mathbb{E}[f_i(\boldsymbol{x})] = \hat{a}_i$ are satisfied. Hence, the problem of finding the maximum entropy distribution is equivalent to optimizing $L(\boldsymbol{\theta})$. We now describe two methods for solving the optimization problem.

##### 1. *Iterative scaling*

Iterative scaling [6] is one of the earliest methods developed for numerically solving the maximum entropy problem. The updating scheme for $\boldsymbol{\theta}$ is the following,

$$\boldsymbol{\theta}_{t+1} = \boldsymbol{\theta}_t + \frac{\gamma}{\sum_i \langle f_i \rangle_{\boldsymbol{\theta}_t}} \ln \frac{\langle \boldsymbol{f} \rangle_{\boldsymbol{\theta}_t}}{\hat{\boldsymbol{a}}} \tag{8}$$

where $\langle \boldsymbol{f} \rangle_{\boldsymbol{\theta}_t}$ is the vector associated with the constraint functions $f_1, f_2, \cdots, f_m$ evaluated at the current $\theta_1, \theta_2, \cdots, \theta_m$, $\gamma$ is the constant learning rate.

##### 2. *Gradient descent (GD)*

A typical procedure to implement the GD method involves solving,

$$\boldsymbol{\theta}_{t+1} = \boldsymbol{\theta}_t - \gamma_t \nabla L(\boldsymbol{\theta}_t) \tag{9}$$

where the step size (learning rate) is either a constant ($\gamma_t = \gamma$) or allowed to change at each step, $t$. The gradient of $L(\boldsymbol{\theta})$ is,

$$\frac{\partial L(\boldsymbol{\theta})}{\partial \theta_i} = \langle f_i \rangle_{\boldsymbol{\theta}} - \hat{a}_i \tag{10}$$

where the subscript $\boldsymbol{\theta}$ indicates the average is evaluated at the parameters $\boldsymbol{\theta}$.

###### *3. Regularization*

For Chromosome 2, investigated in the main text, the positions of ∼900 loci were measured, resulting in $\approx 900 \times 900/2 \approx 4 \times 10^5$ parameters (number of $k_{ij}$ values) that need to be determined. With this many parameters, regularization is usually needed in order to avoid overfitting. In this work, unless specified otherwise, we employ the $L_2$ regularization, which adds a penalty term to the objective function,

$$\tilde{L}(\boldsymbol{\theta}) = \ln Z(\boldsymbol{\theta}) - \boldsymbol{\theta}^T \hat{\boldsymbol{a}} + \lambda ||\boldsymbol{\theta}||_2^2. \tag{11}$$

The penalty term $\lambda ||\boldsymbol{\theta}||_2^2$ constrains the values of $\theta_i$ to be close to zero. The parameter $\lambda$ controls the contribution of the penalty term. To minimize $\tilde{L}(\boldsymbol{\theta})$, we employ the gradient descent method. The gradient of $\tilde{L}(\boldsymbol{\theta})$ is given by,

$$\frac{\partial \tilde{L}(\boldsymbol{\theta})}{\partial \theta_i} = \langle f_i \rangle_{\boldsymbol{\theta}} - \hat{a}_i + 2\lambda\theta_i. \tag{12}$$

##### B. Practical considerations

In DIMES, the constraints are the mean squared pair-wise distances, $\langle r_{ij}^2 \rangle$, where $r_{ij}$ is the distance between loci $i$ and $j$ and $\langle \cdot \rangle$ denotes the ensemble average. The total number of the constraints is $N(N-1)/2$ where $N$ is the total number of loci.

##### 1. *Obtaining the mean squared pair-wise distance matrix*

If the input data is direct measurements of the 3D coordinates of certain loci in the chromatin, then the mean squared pair-wise distances between all pairs are computed as,

$$\langle r_{ij}^2 \rangle = \frac{1}{M} \sum_{m=1}^{m=M} ||\boldsymbol{x}_i^{(m)} - \boldsymbol{x}_j^{(m)}||^2 \tag{13}$$

where $\boldsymbol{x}_i^{(i)}$ and $\boldsymbol{x}_j^{(i)}$ are the 3D coordinates of $i$ and $j$ loci in the $m^{th}$ single cell and $M$ is the total number of cells for which measurements are made.

If the input data is the Hi-C contact frequencies, then the mean squared pair-wise distances between all the pairs are computed using [3],

$$\langle r_{ij}^2 \rangle = \left( \Lambda p_{ij}^{-1/\alpha} \right)^2 \tag{14}$$

where $\Lambda$ sets the length scale and $\alpha$ determines the power-law relation between spatial distances and contact probability. In practice, we choose $\alpha = 4$ if it is not determined experimentally. The value of $\Lambda$ can only be determined when a length scale of the system is known. For instance, if the average radius of gyration of the system is known, $\Lambda$ could be determined by matching the average radius of gyration of the chromosome from the model with the known value. In practice, we simply set $\Lambda = 1$ if not explicitly specified. Note that the structures obtained with $\Lambda = 1$ can be simply rescaled.

##### 2. *Optimization*

Let us denote the target mean squared pair-wise distances as $\langle r_{ij,\text{exp}}^2 \rangle \equiv a_{ij}$. The updating scheme for the values of $k_{ij}$ in the iterative scaling method is,

$$k_{ij}(t+1) = k_{ij}(t) + \frac{\gamma}{\sum_{i<j} \langle r_{ij}^2 \rangle (t)} \ln \frac{\langle r_{ij}^2 \rangle (t)}{a_{ij}} \tag{15}$$

$t$ is the step number, and $\langle r_{ij}^2 \rangle (t)$ is the value of mean squared distance between loci $i$ and $j$ with parameters $k_{ij}(t)$ at step $t$. The value of the constant learning rate, $\gamma$, is chosen to be $\gamma = 10$ because it gives good convergence speed while ensuring that the result converges.

For gradient descent (GD), the updating scheme for $k_{ij}$ is,

$$k_{ij}(t+1) = k_{ij}(t) - \gamma\big[\langle r_{ij}^2\rangle(t) - a_{ij}\big] \tag{16}$$

where $\gamma$ is the constant learning rate.

For both of these two methods, the value of $\langle r_{ij}^2\rangle(t)$ need to be evaluated. In DIMES, the fact that the maximum entropy distribution is a multivariate normal distribution allows us to compute $\langle r_{ij}^2\rangle(t)$ directly using the Eq.4. With values of $k_{ij}(t)$, the matrix $\boldsymbol{K}(t)$ is constructed according to Eq.3. Then, the matrix $\boldsymbol{\Sigma}(t)$ is calulated using $\boldsymbol{\Sigma}(t) = -\boldsymbol{K}^{+}(t)$ where $\boldsymbol{K}^{+}(t)$ is the Moore-Penrose inverse of $\boldsymbol{K}(t)$. Finally, $\langle r_{ij}^2\rangle(t)$ is computed using $\langle r_{ij}^2\rangle(t) = 3\big[\Sigma_{ii}(t) + \Sigma_{jj}(t) - 2\Sigma_{ij}(t)\big]$. For Moore-Penrose inverse, we use Python *SciPy* package [7].

Once the values of $\langle r_{ij}^2\rangle(t)$ are obtained, the values of $k_{ij}$ are updated using Eq.15 for iterative scaling or Eq.16 for GD.

##### 3. *Generation of structures*

After $T$ number of iteration steps, a reasonably converged $\boldsymbol{K}(T)$ is obtained. Denote $\boldsymbol{K} \equiv \boldsymbol{K}(T)$. To generate an ensemble of structures, we sample the distribution given by Eq.2. Following the derivation in our previous work [1], the coordinates of loci, $\boldsymbol{X}_p$ ($p$ denotes the three spatial dimensions), can be computed as a linear combination of normal modes, $\boldsymbol{X}_p = \boldsymbol{V}^T\boldsymbol{R}_p$ where $\boldsymbol{R}_p$ are the normal modes. $\boldsymbol{V}$ is obtained by eigendecomposition of $\boldsymbol{K}$, i.e. $\boldsymbol{V}\boldsymbol{K}\boldsymbol{V}^T = \boldsymbol{\Omega} = \mathrm{diag}(\omega_1, \omega_2, \cdots, \omega_N)$. Each component of the normal modes, $R_{i,p}$, is a Gaussian random variable with distribution $\mathcal{N}(0, -\omega_i^{-1})$. Note that different $R_{i,p}$ are independent to each other.

The procedures used to generate an ensemble of structures involve the following steps:

(a) Perform eigendecomposition of $\boldsymbol{K}$. Obtain $\boldsymbol{V}$ and the eigenvalues $\omega_1, \omega_2, \cdots, \omega_N$.

(b) Draw a total of $N$ random numbers from distributions $\mathcal{N}(0, -\omega_i^{-1})$ for $i = 1, 2, \cdots, N$. Denote these random numbers as $R_1, R_2, \cdots, R_N$.

(c) Define $\boldsymbol{R} = [R_1, R_2, \cdots, R_N]^T$. Compute $\boldsymbol{X}$ using $\boldsymbol{X} = \boldsymbol{V}^T\boldsymbol{R}$.

(d) Repeat (b) and (c) two more times, each for a spatial dimension. Finally, we obtain $\boldsymbol{X}_1, \boldsymbol{X}_2, \boldsymbol{X}_3$, representing the $x$, $y$ and $z$ coordinates of $N$ number of loci.

(e) Repeat (b) - (d) $M$ times, resulting a total number of $M$ randomly sampled 3D structures.

#### SUPPLEMENTARY NOTE 4 – CORRELATION MATRIX

In this note, we describe how to calculate the correlation matrix given the connectivity matrix $\boldsymbol{K}$. Let us denote the correlation matrix as $\boldsymbol{\rho}$. The elements of $\boldsymbol{\rho}$, $\rho_{ij}$, are defined as,

$$\rho_{ij} = r_s(\boldsymbol{K}_i, \boldsymbol{K}_j) \tag{17}$$

where $\boldsymbol{K}_i$ and $\boldsymbol{K}_j$ are the $i^{th}$ row and $j^{th}$ row of matrix $\boldsymbol{K}$, respectively, $r_s(\cdot)$ is the standard function used to compute the Spearman's rank correlation coefficient between $\boldsymbol{K}_i$ and $\boldsymbol{K}_j$. The calculation is done using *SciPy* Python package [7]

#### SUPPLEMENTARY NOTE 5 – STRUCTURAL VARIATIONS

In this note, we describe how the connectivity matrix $\boldsymbol{K}$ is modified in modeling of structural variations (shown schematically in Fig.5), which are mutations in the chromosomes. Our goal is to utilize the calculated $\boldsymbol{K}$ for the wild-type chromosome, and then predict the structural changes in the chromosome *without* any additional inputs or modifications.

##### Inversion

Inversion means that a genome sequence is reversed from end-to-end. Let the inverted segment lie between $m^{th}$ and $n^{th}$ loci ($m < n$) in the wild type (WT). We account for the effects of inversion is by modifying only the elements of $\boldsymbol{K}$ associated with the loci involved in the inversion process. More precisely, the modification of $\boldsymbol{K}$ can be summarized as,

$$
\begin{aligned}
K_{ij}^{\mathrm{INV}} &= K_{ij}^{\mathrm{WT}}, \text{ if } i \not\subset [m,n] \text{ and } j \not\subset [m,n] \\
K_{ij}^{\mathrm{INV}} &= K_{i,m+n-j}^{\mathrm{WT}}, \text{ if } i < m \text{ and } j \subset [m,n] \\
K_{ij}^{\mathrm{INV}} &= K_{m+n-i,j}^{\mathrm{WT}}, \text{ if } j > n \text{ and } i \subset [m,n] \\
K_{ij}^{\mathrm{INV}} &= K_{m+n-i,m+n-j}^{\mathrm{WT}}, \text{ if } i \subset [m,n] \text{ and } j \subset [m,n]
\end{aligned}
$$

where the superscripts denote inversion (INV) and wild type (WT), respectively.

**Deletion**

Deletion corresponds to a deletion of a genome sequence, which would shorten the number of base pairs. As before, let the segment deleted be located between $m^{th}$ and $n^{th}$ loci ($m < n$) in the wild type. The deletion effect is modeled by removing the $k_{ij}$ elements from $\boldsymbol{K}$ involving the deleted sequence. The modification of $\boldsymbol{K}$ is implemented using,

$$
\begin{aligned}
K_{ij}^{\mathrm{DEL}} &= K_{ij}^{\mathrm{WT}}, \text{ if } i,j < m \text{ or } i,j > m \\
K_{ij}^{\mathrm{DEL}} &= K_{i,n+j-m}^{\mathrm{WT}}, \text{ if } i < m \text{ and } j > m \\
K_{ij}^{\mathrm{DEL}} &= (K_{m-1,m}^{\mathrm{WT}} + K_{m,m+1}^{\mathrm{WT}})/2, \text{ if } i = m-1 \text{ and } j = m
\end{aligned}
$$

where the superscript DEL denotes deletion.

In both inversion and deletion variants, we set $K_{ij} = K_{ji}$. The diagonal elements in $\boldsymbol{K}$ is computed using $K_{ii} = -\sum_{j \neq i} K_{ij}$. It is worth emphasizing that in determining the effects of inversion and deletion the $\boldsymbol{K}$ matrix is determined only once for the wild type.

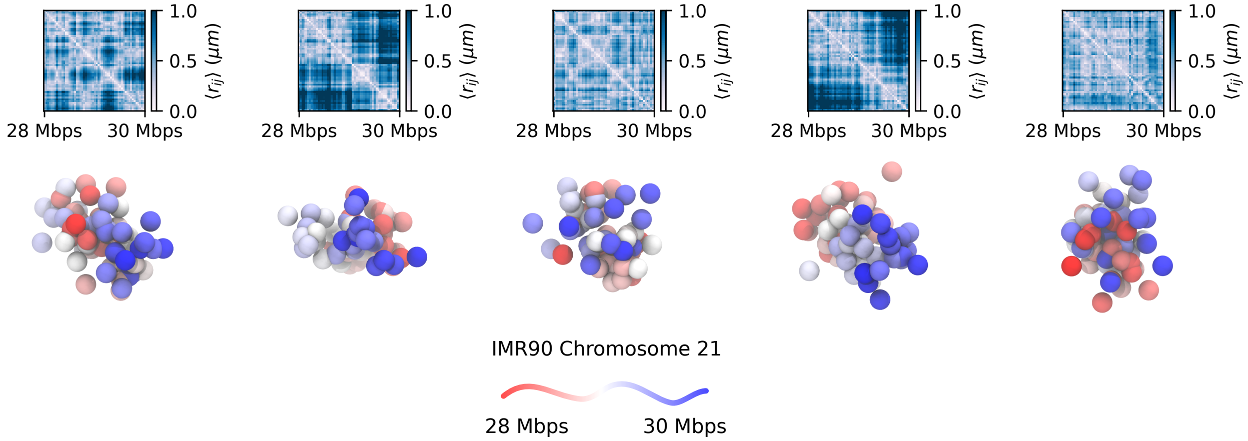


FIG. 1. Examples of the distance map for a few individual conformations for Chr21 in the 28 Mbps - 30 Mbps region in IMR90 cell line. Each panel shows an individual structure randomly chosen from the ensemble from which the spatial distance matrix is computed. The distance scale is given on the right of the panels. Just as in experiments [8], TAD-like or domain-like patterns are clearly observed in the spatial distance matrices even at the single cell level. The snapshots below show the structures. Each bead in the structure represents a single locus. The color of the loci in the 3D structures represents the genomic location, ranging from red to blue.

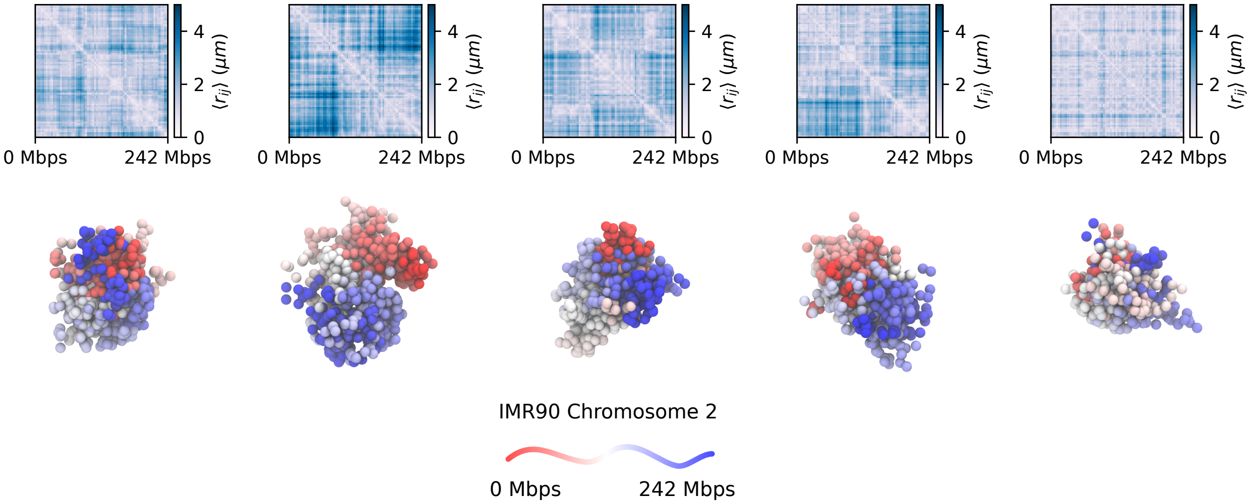


FIG. 2. Distance maps (scales on the right) for individual conformations for the nearly 242 Mbps Chr2 in IMR90 cell line. We find substantial variations in the distance maps in individual conformations, attesting to the heterogenity in the genome organization. This observation accords well with experiments [9]. In the snap shots below, each bead represents one locus. The color of loci in the 3D structures represents the its genomic location, ranging from red to blue.

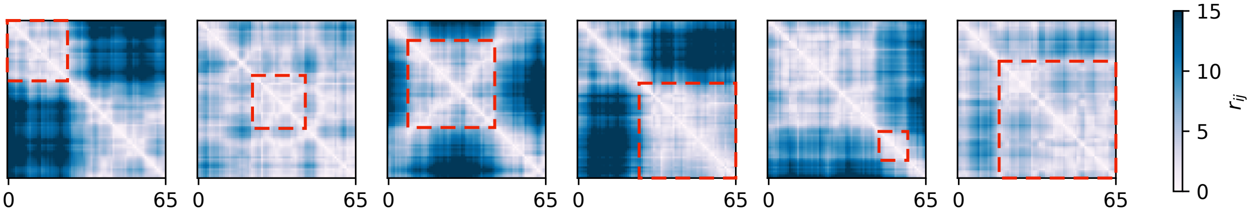


FIG. 3. Distance maps (scales on the right) for random sampled conformations for an ideal Rouse homopolymer (the number of monomers is 65). We find substantial variations in the distance maps in individual conformations, which stem from the intrinsic thermal fluctuations of the ensemble. TAD-like domains are discernible in the distance matrices. A few examples of such TAD-like structures are marked by dashed squares.

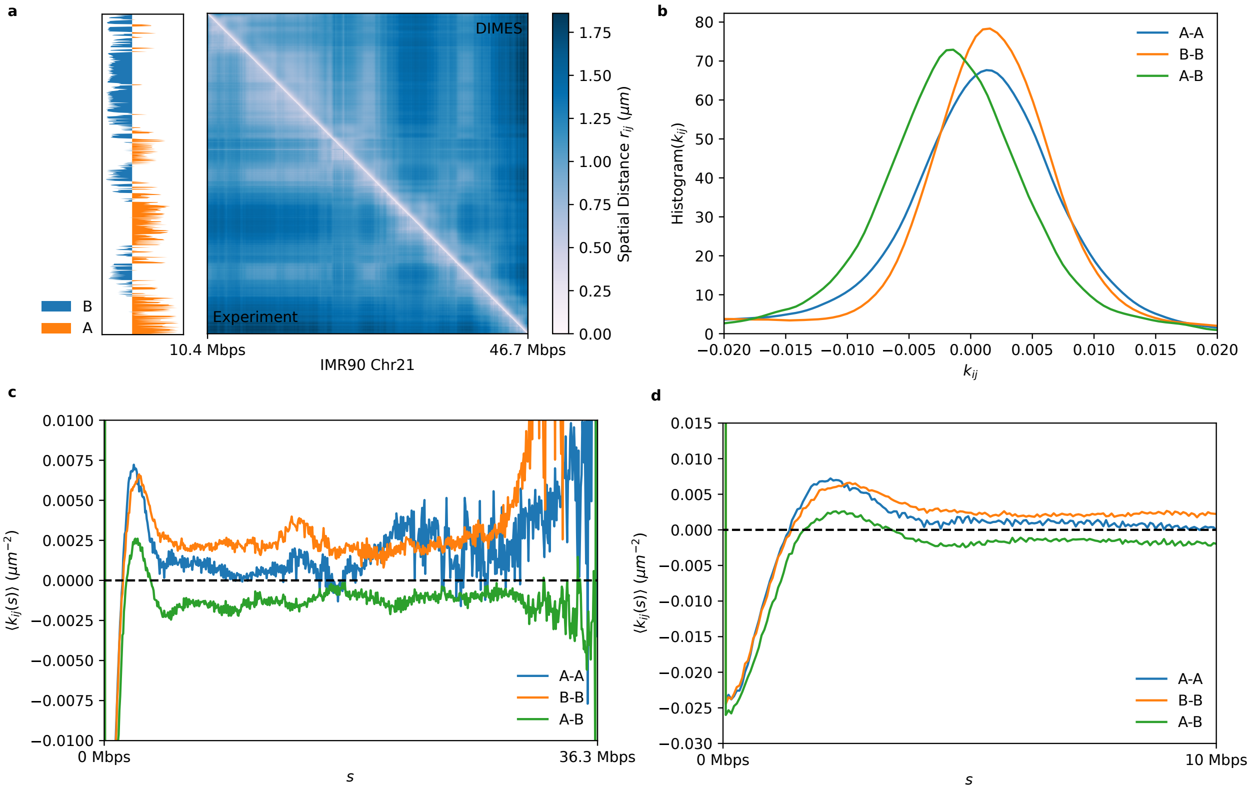


FIG. 4. Characteristics of pair-wise interactions of the Chromosome 21 for IMR90 cell line. **(a)** The average distance matrix between experiment (lower triangle) and the DIMES (upper triangle) show excellent agreement. The left track shows the principle component dimension 1 (PC1) computed using principle component analysis (PCA) from the connectivity matrix $\boldsymbol{K}$. **(b)** Histogram of $k_{ij}$ for A-A, B-B and A-B. **(c)** Genomic-distance normalized $\langle k_{ij}(s) \rangle = (1/(N-s)) \sum_{i<j}^{N} \delta(s-(j-i))k_{ij}$ for A-A, B-B and A-B. $\langle k_{ij}(s) \rangle$ are shown for $s$ between 0 and 36.3 Mbps. **(d)** Enlarge portion of **(c)**, showing $\langle k_{ij}(s) \rangle$ for $s < 10$ Mbps.

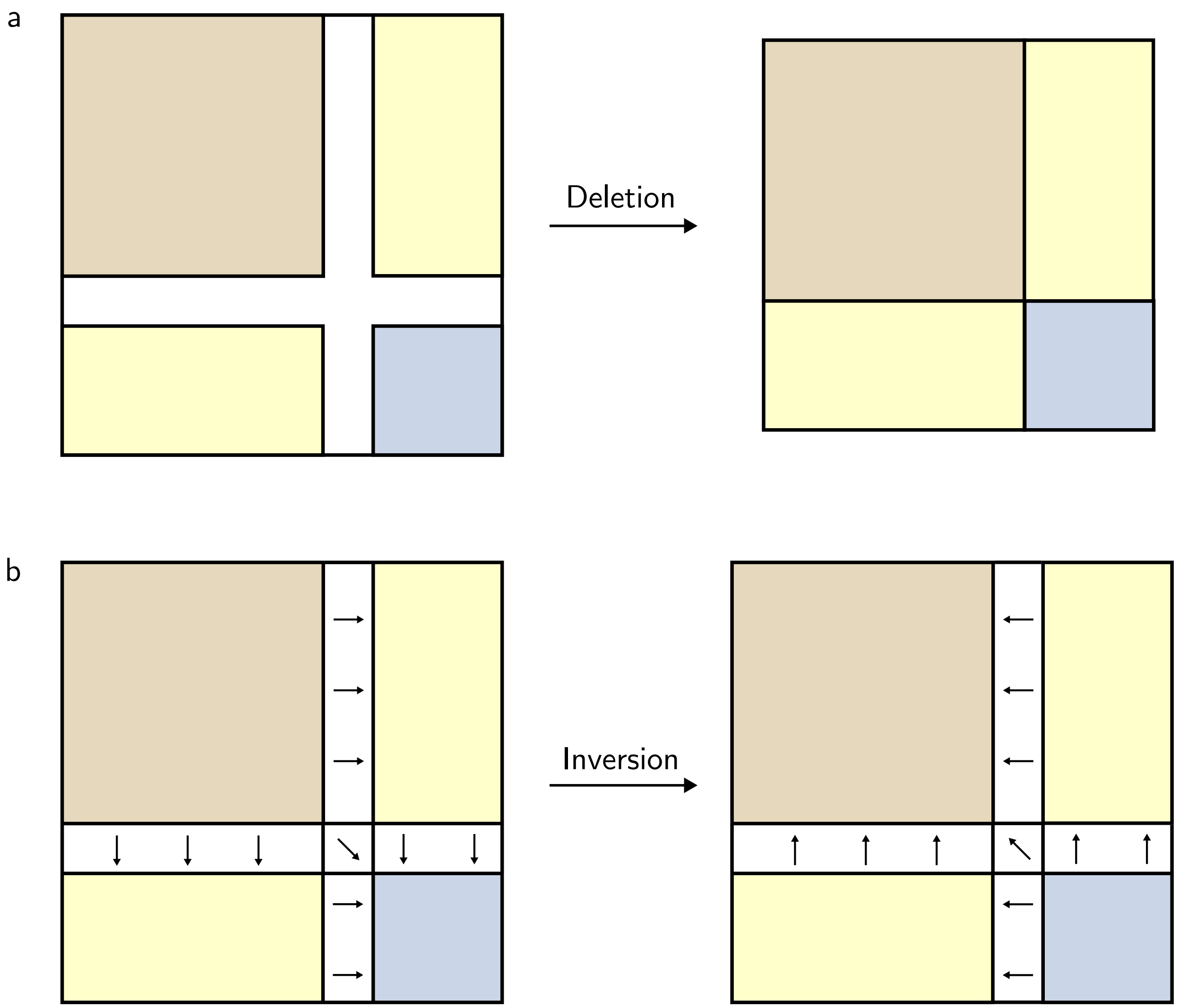


FIG. 5. Schematics for deletion operation and inversion operation on connectivity matrix $\boldsymbol{K}$. **(a)** Illustration of deletion. Different parts of the matrix $\boldsymbol{K}$ are in different colors. The gray area corresponds to the region deleted. The resulting $\boldsymbol{K}$ is shown on the right. **(b)** The gray areas with gradient correspond to the inverted segment. The change of direction of gradient after the inversion depict how the values of elements in $\boldsymbol{K}$ are altered.

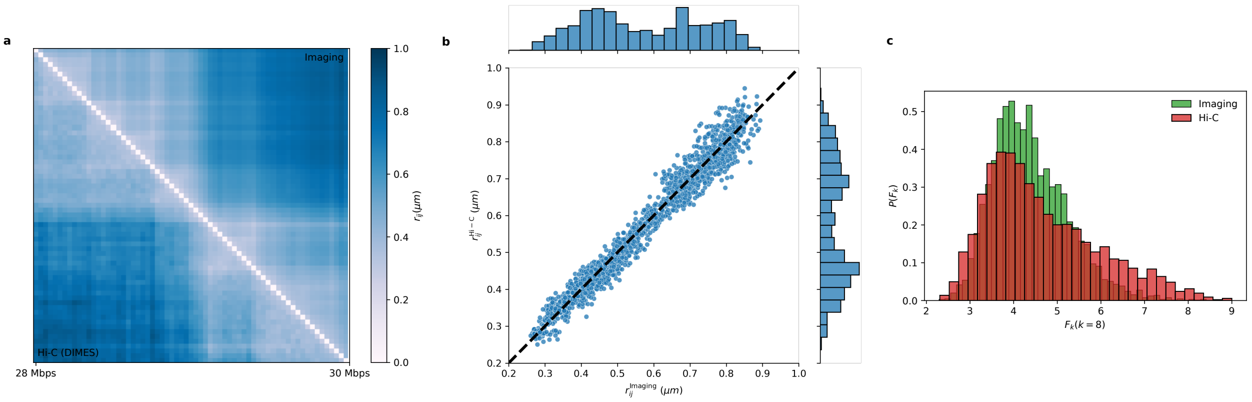


FIG. 6. Structural organization calculated from Hi-C and imaging data for Chr21 28 Mbps - 30 Mbps from IMR90 cell line. **(a)** Comparison between the mean distance matrix inferred from Hi-C contact map (lower triangle) and the experimental measured average distance matrix (upper triangle). The distance scale is shown on the right. **(b)** Direct comparison of pairwise distances, $\langle r_{ij}^{\text{Imaging}} \rangle$ versus $\langle r_{ij}^{\text{Hi-C}} \rangle$. Each dot represents a pair $(i, j)$. Dashed line, with a slope of unity, is a guide to the eye. **(c)**. Histogram of $F_k(k = 8)$ for 4,000 individual structures.

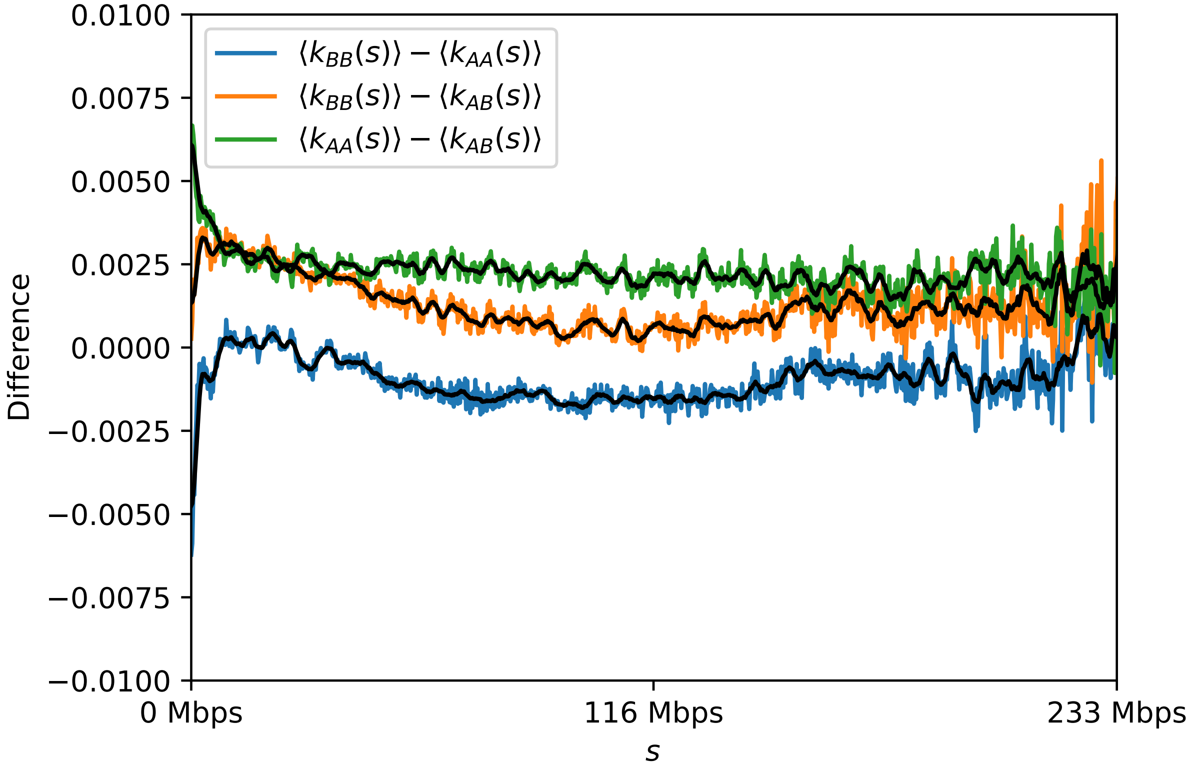


FIG. 7. $\langle k_{AA}(s) \rangle - \langle k_{AB}(s) \rangle$, $\langle k_{AA}(s) \rangle - \langle k_{BB}(s) \rangle$, and $\langle k_{BB}(s) \rangle - \langle k_{AB}(s) \rangle$ as a function of the genomic distance $s$. Black lines are the moving average over window size of $s = 2.6$Mbps

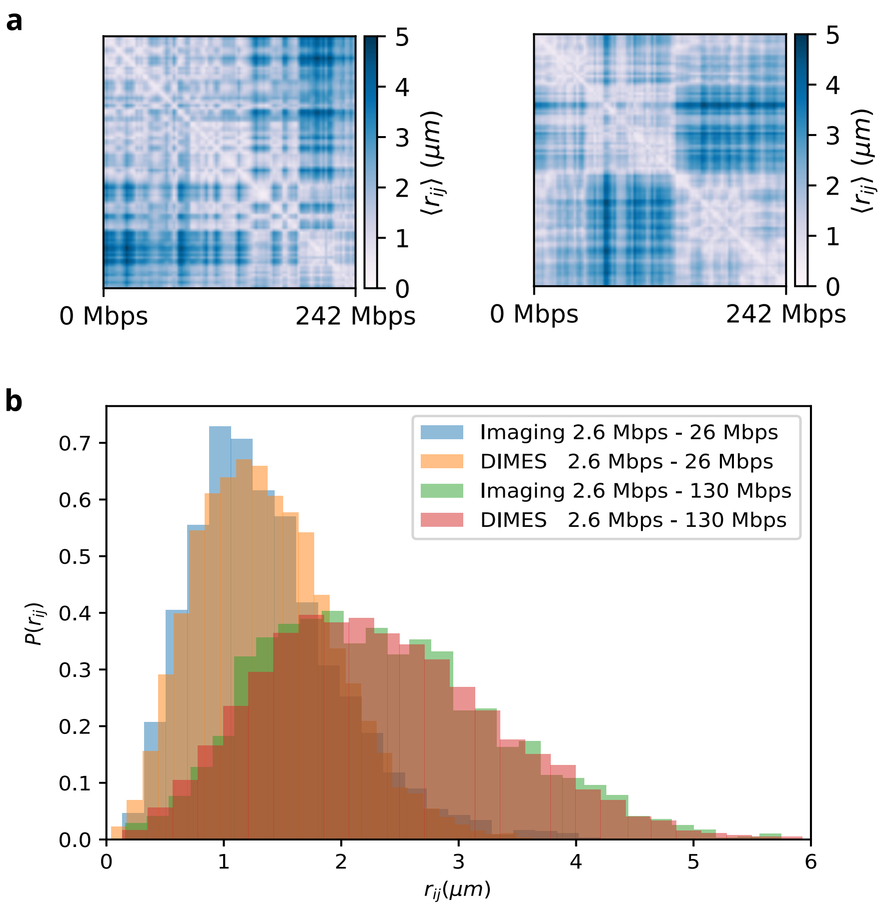


FIG. 8. **(a)** Experimental single-cell distance maps for Chr2 calculated from imaging data [9]. **(b)** Comparison of distributions of pairwise distances between imaging data and the prediction from DIMES. Pair 2.6 Mbps - 26 Mbps and pair 2.6 Mbps - 130 Mbps are shown.

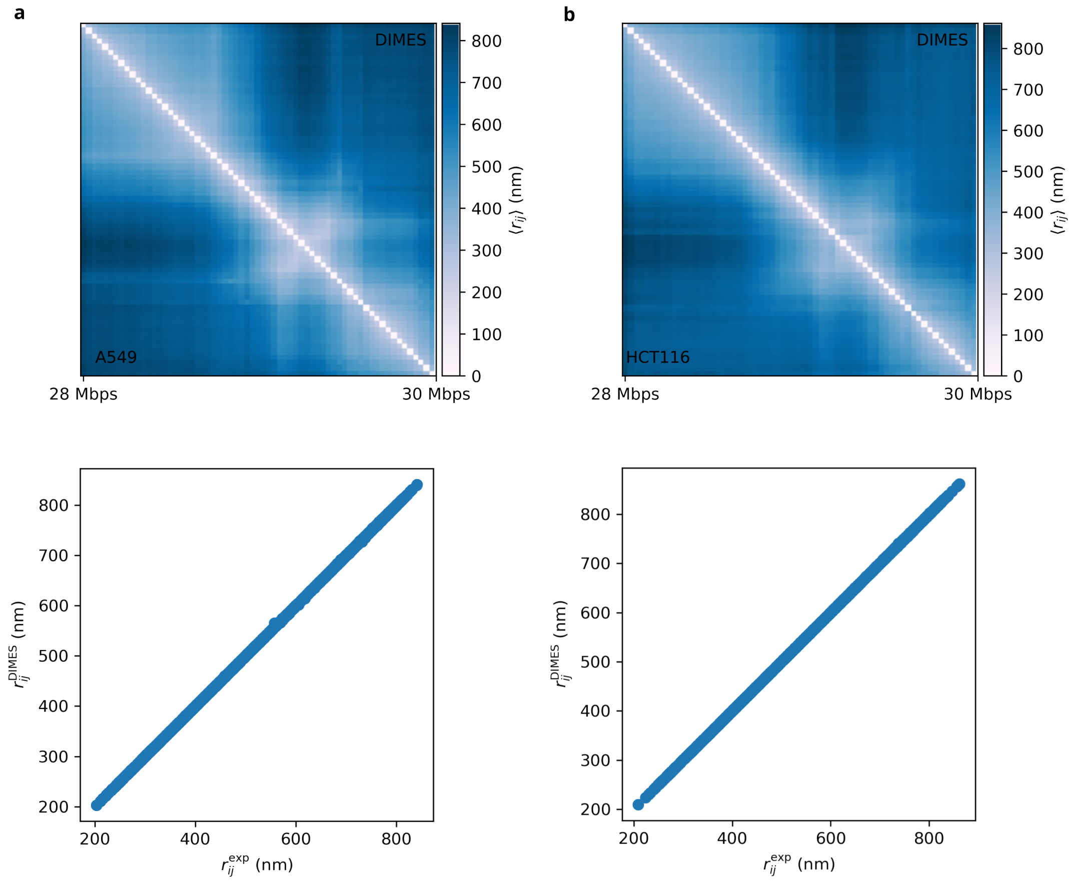


FIG. 9. Efficacy of DIMES in matching the targets generated using experimental imaging data. Comparison between the mean spatial distances computed from the reconstructed structures and the experimental data in cell lines: A549 (**a**), and HCT116 (**b**). The upper panel shows side-by-side comparisons of the distance matrices, and the lower panel displays the scatter plot between individual pairwise distances $r_{ij}$'s.

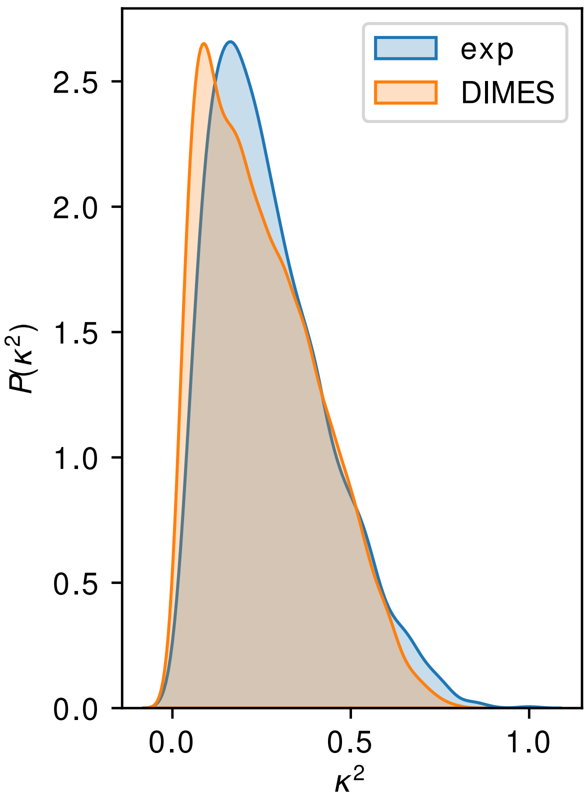


FIG. 10. Comparison of $P(\kappa^2)$ between experiment and the DIMES predictions. $P(\kappa^2)$ is the probability density distribution of the shape parameter $\kappa^2$ for the 28 Mbps - 30 Mbps region of Chr21.
